# Coupling between lipid miscibility and phosphotyrosine driven protein condensation at the membrane

**DOI:** 10.1101/2020.07.22.215970

**Authors:** J. K. Chung, W. Y. C. Huang, C. B. Carbone, L. M. Nocka, A. N. Parikh, R. D. Vale, J. T. Groves

## Abstract

Lipid miscibility phase separation has long been considered to be a central element of cell membrane organization. More recently, protein condensation phase transitions, into three-dimensional droplets or in two-dimensional lattices on membrane surfaces, have emerged as another important organizational principle within cells. Here, we reconstitute the LAT:Grb2:SOS protein condensation on the surface of giant unilamellar vesicles capable of undergoing lipid phase separations. Our results indicate that assembly of the protein condensate on the membrane surface can drive lipid phase separation. This phase transition occurs isothermally and is governed by tyrosine phosphorylation on LAT. Furthermore, we observe that the induced lipid phase separation drives localization of the SOS substrate, K-Ras, into the LAT:Grb2:SOS protein condensate.

**Statement of Significance:** Protein condensation phase transitions are emerging as an important organizing principles in cells. One such condensate plays a key role in T cell receptor signaling. Immediately after receptor activation, multivalent phosphorylation of the adaptor protein LAT at the plasma membrane leads to networked assembly of a number of signaling proteins into a two-dimensional condensate on the membrane surface. In this study, we demonstrate that LAT condensates in reconstituted vesicles are sufficient to drive lipid phase separation. This lipid reorganization drives another key downstream signaling molecule, Ras, into the LAT condensates. These results show that the LAT condensation phase transition, which is actively controlled by phosphorylation reactions, extends its influence to control lipid phase separation in the underlying membrane.

## Introduction

In 1973, shortly after the classic fluid mosaic description of cell membranes was published (1), a series of papers from Harden McConnell’s lab described discovery of lateral phase separation in the lipids of cell membranes (2–6). Contemporary work from Erich Sackmann and colleagues confirmed an intriguing heterogeneity in the organization of lipids in the fluid membrane (7). This phenomenon later developed into the lipid raft model of cell membranes, as articulated by Kai Simons and Elina Ikonen in 1997 (8–10). The field of lipid rafts has since both flourished and attracted great controversy (11–14). Although lipid miscibility phase transitions are readily, and spectacularly visualized in purified lipid membranes (15–17), their unambiguous detection in living cells proved much more challenging (18–21), with only very limited definitive observations reported (22). There is evidence that cell membranes are poised near a miscibility phase transition (23), which naturally leads one to speculate that this may be actively controlled by the cell. However, a longstanding criticism of the lipid raft model questions how lipid phase separation could be controlled with the specificity required for biological functions, while the underlying interactions between lipids and cholesterol that enable the phase transition are rather nonspecific (24). Clearly proteins must play a commanding role controlling lipid phase separation in the physiological setting, but we have very limited mechanistic understanding of how this is actually achieved in specific cases (25).

One prominent example that captures this debate is the T-cell receptor (TCR) signaling system. TCR and a number of downstream proteins including linker for activation of T cells (LAT), phospholipase C gamma 1 (PLCγ1), and the Ras activator son of sevenless (SOS), form clusters on the membrane (26–32). Earlier studies using detergent-resistant membrane (DRM) extraction have suggested that these molecules reside on lipid rafts (33–35). However, subsequent studies have failed to conclusively establish lipid rafts as the driving force for TCR-induced signaling clusters (36–38). Furthermore, it remains unclear how signaling activity—in the case of TCR, the receptor activation and tyrosine phosphorylation of downstream proteins including LAT—could trigger the lipid phase separation. This disconnect is further underscored by the fact that, at physiological ligand densities (39), individual TCR are capable of triggering the entire signaling pathway without ever forming clusters themselves (40–44).

Modular binding interactions among proteins present another type of mediated molecular assembly process in cells (45, 46). With sufficient multivalency, these interactions can lead to protein condensation phase transitions into three-dimensional droplets (47), sometimes called membraneless organelles, or two-dimensional assemblies on the membrane surface (48–51). Similar biomolecular condensates can also incorporate nucleic acids and play a role in transcription regulation (52, 53).

It has recently been discovered that LAT can participate in a protein condensation phase transition in reconstituted membranes (48, 49, 54, 55). LAT is a transmembrane scaffold protein that becomes phosphorylated at multiple tyrosines upon TCR activation. Three of the phosphotyrosines on LAT are canonical docking sites for the SH2 domain of growth factor receptor-bound protein 2 (Grb2), a cytosolic adapter protein (56). Grb2 additionally has SH3 domains, which bind to the proline-rich domain of SOS, a guanine nucleotide exchange factor that activates Ras (57). A single SOS can associate with at least two Grb2 molecules, and these multivalent interactions result in an extended two-dimensional network assembly of LAT:Grb2:SOS on the membrane in a phosphorylation-dependent manner (58, 59). This complex has been shown to play an important role in T-cell signaling (60, 61). The LAT:Grb2:SOS protein condensation phase transition is reversible, and since it is governed by tyrosine phosphorylation, it is directly under the control of competing kinase and phosphatase reactions in the TCR signaling system.

Here we reconstitute the LAT:Grb2:SOS protein condensate from purified proteins on giant unilamellar vesicle (GUV) membranes that can undergo lipid phase separation. The cytoplasmic domain of LAT was purified with an N-terminal His_6_ tag and labeled with Alexa Fluor 555 (AF555) at Cys146 via maleimide-thiol chemistry. LAT was phosphorylated by the kinase domain of Hck in solution. Then, phosphorylated LAT (*p*LAT) was linked to the membrane by the binding of His_6_ tag to the Ni-NTA lipids in the membrane. This membrane-linked *p*LAT exhibits free lateral diffusion and remains monomeric prior to any assembly (48, 49). The addition of full-length Grb2 and the proline-rich domains of SOS leads to the networked condensation of LAT:Grb2:SOS on the membrane surface of GUVs, as shown in Figure 2 (top row). Here, the condensates are visualized as concentrated regions of *p*LAT-AF555 fluorescence on GUVs by confocal microscopy. This condensate is mediated by tyrosine phosphorylation on LAT, and is reversible (Figure 2, bottom row). The rapid (<10 s) dispersion of the condensed structure upon phosphatase (YopH) addition indicates that the individual Grb2:pLAT bonds must be highly dynamic and offer little protection from solution phosphatases. Incidentally, the membrane-linked phosphatase CD45 has been reported to be excluded from LAT condensates, possibly providing some degree of positive feedback with respect to this phosphatase (48).

We next examined how the lipid phase transition behavior of GUVs is perturbed by the LAT condensate. GUVs composed of a ternary mixture of saturated lipids, unsaturated lipids, and sterols (in a roughly 1:1:1 ratio) exhibit temperature-dependent miscibility phase separation. Below the miscibility transition temperature, *T*_misc_, the vesicles separate into coexisting liquid ordered (*L*_o_) and liquid disordered (*L*_d_) regions (16, 62). As a crude guideline, the *L*_o_ region is rich in saturated phosphatidylcholine (PC) lipids such as DPPC, while *L*_d_ is rich in unsaturated phospholipids such as DOPC (63). For our experiments, GUVs composed of 29.2% DOPC, 33.2% DPPC, 33.3% cholesterol, 4% Ni-DOGS, 0.1% Texas Red-DHPE (TR-DHPE), and 0.2% Oregon Green-DHPE (OG-DHPE) were used. This composition is an approximation of the well-characterized equimolar mixture of DOPC, DPPC, cholesterol, and the observed *T*_misc_ is also close to the reported value, 29°C (62). Even though it is not critical into which lipid phase LAT partitions in our experiments, the full length protein has been shown to partition into clusters without lipid raft makers (GPI anchors) in live cells (38) – suggesting that LAT does not partition into *L*_o_-like phase in cells. In our experiments, because Ni-DOGS chain is unsaturated (18:1-18:1), Ni-chelated *p*LAT is expected partition into the *L*_d_ region. This is confirmed by its colocalization with TR-DHPE, which is a well-established reporter of *L*_d_ phase (16). On the other hand, TR and OG fluorescence exclude each other upon phase separation, indicating that OG-DHPE partitions into the *L*_o_ phase (Figure 1A).

**Figure 1.**
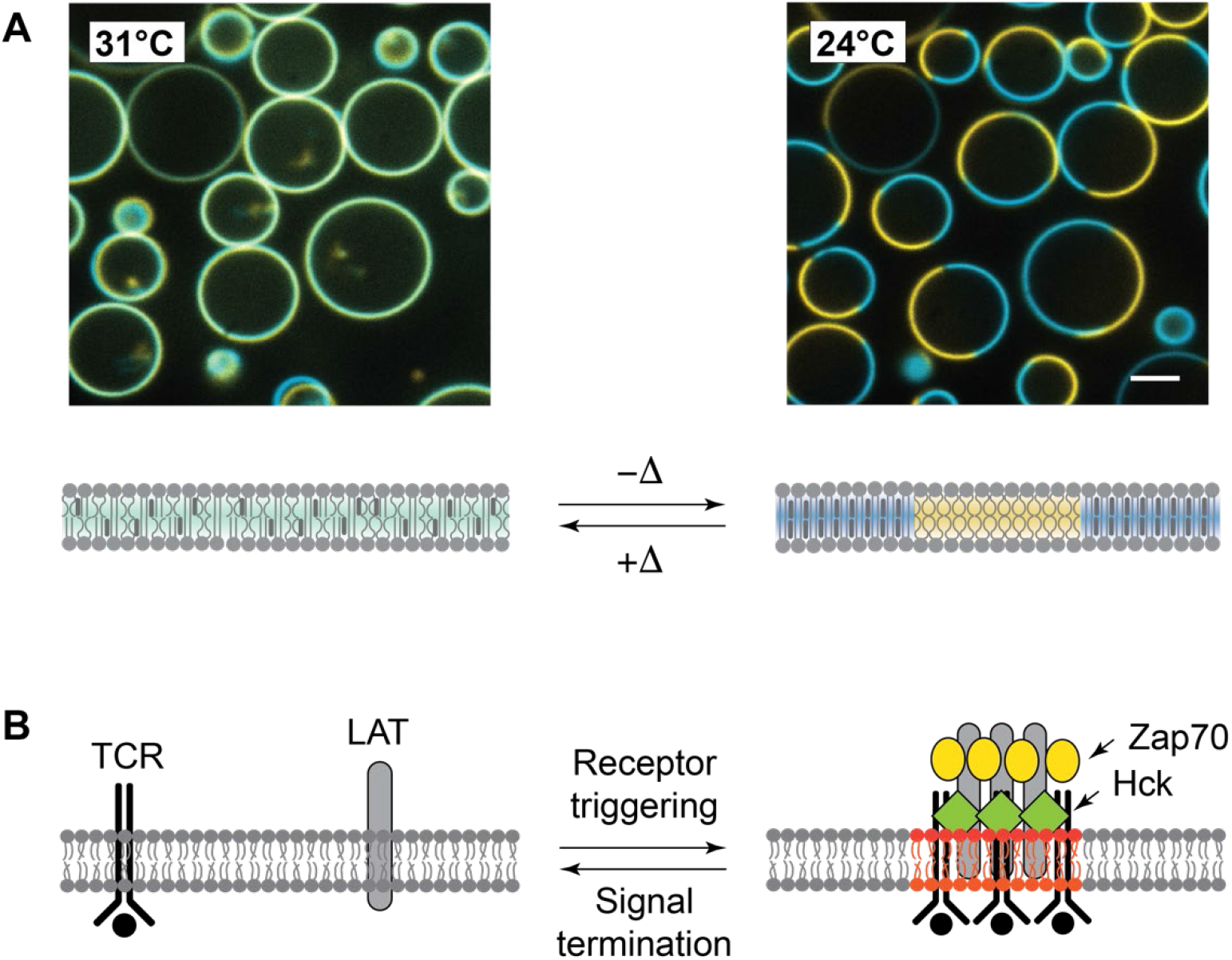
**(A)** Temperature-dependent liquid-liquid phase separation in giant unilamellar vesicles (GUVs). Above the transition temperature *T*_misc_, the distribution of lipids is homogeneous across the membrane (left). Below *T*_misc_, lipids compartmentalize into macroscopic domains: the *L*_d_ domain (yellow, TR-DHPE) is enriched with unsaturated lipids, and the *L*_o_ with saturated lipids (blue, OG-DHPE). **(B)** In lipid raft theory, clusters of signaling proteins such as the T-cell receptors, are “carried” on ordered lipid domains to facilitate signal transduction.

**Figure 2.**
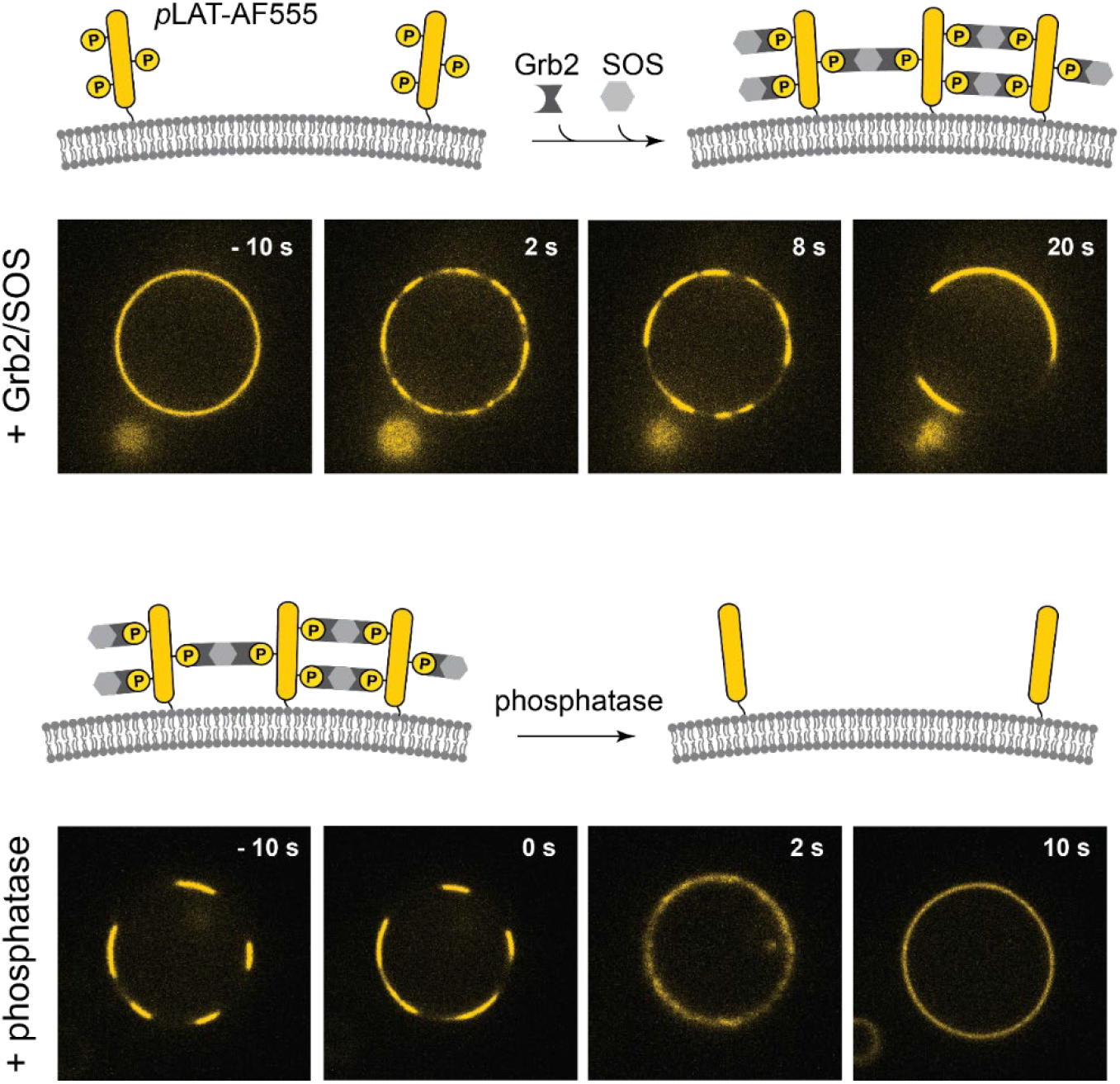
The LAT:Grb2:SOS protein condensate was reconstituted on GUVs. His-tagged phosphor-LAT (*p*LAT) is associated with the vesicles by chelating to Ni-NTA lipids. The introduction of Grb2 and SOS results in extended networks of LAT condensate, visualized by the AF555 fluorescence in the confocal microscopy (top). The LAT:Grb2:SOS assembly can be reversed by dephosphorylation of LAT by phosphatase (YopH) (bottom).

First, we examined whether LAT condensation could induce phase transitions in initially uniform vesicles near the miscibility transition temperature. The experiment is shown in Figure 3. In the imaging chamber maintained at 31°C, the *p*LAT-associated vesicle membranes exhibit a homogeneous distribution of fluorescent markers (TR-DHPE and OG-DHPE), as expected since this temperature is slightly above the *T*_misc_ of 29°C. The addition of Grb2-AF647 and SOS triggers a rapid LAT:Grb2:SOS condensation on the membrane surface, which is readily visualized by the appearance of concentrated regions of 647 nm fluorescence, tracking Grb2. This is accompanied by a clear partitioning of TR-DHPE (yellow) and OG-DHPE (blue), indicating a miscibility phase transition within the lipids has also occurred, although here under isothermal conditions. The LAT condensate is coincident with the *L*_o_ region marked by TR-DHPE, while *L*_d_ region, visualized by OG-DHPE, is excluded from LAT.

**Figure 3.**
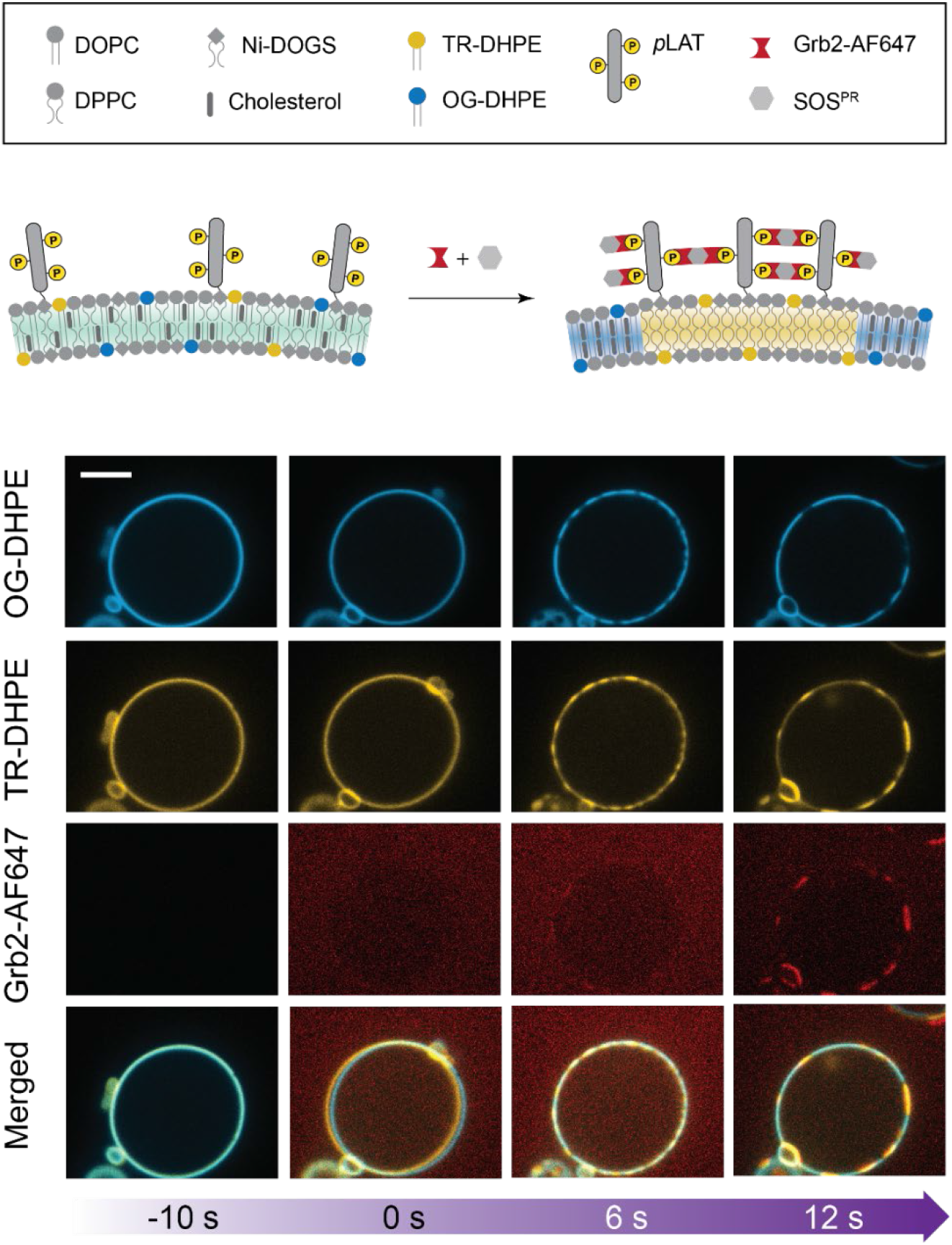
The LAT:Grb2:SOS condensate induces lipid phase separation on GUVs. Starting with a temperature (31°C) above its *T*_misc_ (29°C), the lipids is spatially homogeneous initially. As the proteins assembled (visualized by Grb2-AF647), the lipids undergo liquid-liquid phase transition. OG-DHPE and TR-DHPE mark the *L*_o_ and *L*_d_ regions, respectively.

Next, we examined the temperature-dependent phase separation behavior of the vesicles in equilibrium under different conditions (Figure 4A). In this experiment, the temperature of the imaging chamber was increased gradually, at a rate of about 1°C/min, and held at each temperature data point for 2 min. Then, multichannel confocal images of a population (*n* ~100) of vesicles were obtained. The chamber was cooled back to 20°C at the same rate. The number of phase-separated vesicles were counted at each temperature point. All observations were the same regardless of the direction of the temperature ramp, indicating that all processes, including protein assembly and lipid phase separation, are reversible. Bare GUVs (empty black circles) show *T*_misc_ of 29°C. With *p*LAT associated with the vesicles (solid black circles), *T*_misc_ is shifted slightly but remains essentially the same at 28°C. When the LAT condensate is formed by the addition of Grb2 and SOS, however, the apparent *T*_misc_ is increased to 34°C (red solid circles). This is consistent with the previous experiment in which the lipid phase separation is driven by the protein condensate at a temperature where it would otherwise be homogeneous.

**Figure 4.**
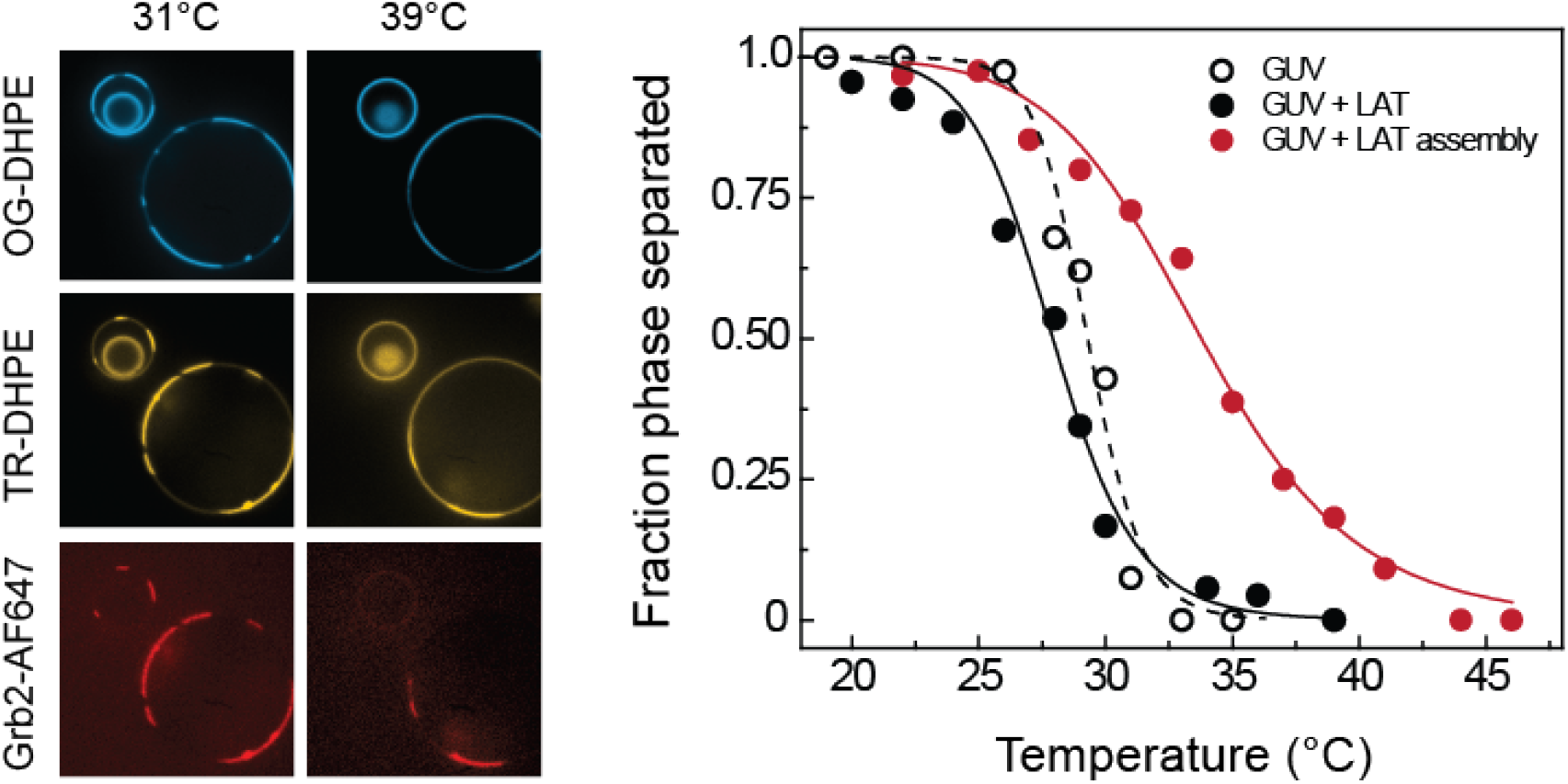
(Left) At 31°C, the GUVs associated with LAT:Grb2:SOS clusters are phase-separated. Note that the smallest vesicle, invaginated within a larger vesicle, is inaccessible to the proteins, and remains homogeneous. At 39°C, Grb2 has dissociated from the smaller vesicle, which became homogeneous. For the larger vesicle on which the protein condensate remains, lipid the phase separation also remains. (Right) The miscibility transition temperatures were measured for bare GUVs, GUVs with LAT, and GUVs with the LAT condensate. The difference in Tm between bare GUVs and LAT-associated GUVs are minimal. However, it is increased significantly in presence of the protein assembly. The data primarily reflect temperature-dependent LAT:Grb2:SOS interactions rather than GUV phase separation, as the protein assembly becomes unstable at high temperatures and dissociate from the vesicles. However, hypothetically, stable LAT:Grb2:SOS interactions would further increase the apparent *T*_misc_.

The apparent Δ*T*_misc_ of 5°C in the presence of the protein condensate is not actually a shift in the lipid *T*_misc_. Rather, the protein condensate itself becomes unstable at higher temperatures, and Grb2 and SOS are released from the vesicle surfaces. This can be seen in Figure 4A, bottom right: at 39°C, the Grb2-AF647 fluorescence is not redistributed on the membranes, but rather reduced overall because it was lost to the solution. The fluorescence signal is recovered when the temperature is lowered, indicating that the protein condensation is also a reversible, temperature-dependent process. As long as the protein condensate is present, vesicles remained phase-separated with the *L*_d_ region templating the protein condensate (Figure 4). This suggests that the actual Δ*T*_misc_ is greater than the apparent value of 5°C, and probably lies outside the experimentally accessible temperature range where both GUV phase separation and LAT:Grb2:SOS condensate can be observed.

Finally, we investigated how the spatial organization of other membrane-bound proteins might be directed by the protein assembly-induced lipid phase separation. K-Ras is a small GTPase and a substrate of SOS, and SOS-catalyzed nucleotide exchange from its GDP-to its GTP-bound state triggers downstream signal activation. The various Ras isoforms serve as hubs for signaling pathways such as phosphoinositide 3-kinases (PI3K) and mitogen-activated protein kinase (MAPK), and Ras misregulation is among the most common causes of cancer (64, 65). Native K-Ras is localized to the membrane by a farnesyl lipid modification, as well as electrostatic interactions between its positively charged region and anionic phospholipids in cellular membranes (66). Therefore, the organization of lipids is expected to play an important role in determining the location of K-Ras. Previous studies have shown that K-Ras partitions to the *L*_d_ region on GUVs, largely due to the highly branched farnesyl anchor (67). Therefore we anticipated K-Ras may similarly be directed by the lipid phase separation induced by the LAT:Grb2:SOS condensate. To examine this, eGFP-labeled full-length K-Ras with its native membrane anchor, including both the farnesylation and methylation of the terminal cysteine (68, 69) (20 nM final concentration), was introduced into GUVs of similar composition as in the previous experiments, but with 10% anionic DOPS lipids (final composition: 19.3% DOPC, 33.3% DPPC, 33.3% cholesterol, 10% DOPS, 4% Ni-DOGS, 0.1% TR-DHPE). The negatively charged lipids are necessary for the stable association of K-Ras to the membrane (69–71). The bottom panel of Figure 5 shows that, prior the introduction of Grb2 and SOS, eGFP-K-Ras (blue) as well as TR-DHPE (yellow) are initially distributed homogeneously on the vesicles. After Grb2 (red) and SOS are added, the lipid membrane becomes phase-separated as the protein assemblies form on its surface. K-Ras, LAT:Grb2:SOS, and TR are observed to partition together in the *L*_d_ region. Coupling of the lipid miscibility phase separation to the LAT:Grb2:SOS protein condensation localizes K-Ras with the condensate. As K-Ras does not colocalize with the protein condensate on supported lipid bilayers that are incapable of phase transitions (Figure S1), its partitioning on GUVs is likely to be driven by its anchor participating in the lipid phase transition. This phase transition and subsequent protein colocalization between SOS and K-Ras occurs isothermally, and under the control of tyrosine phosphorylation reactions.

**Figure 5.**
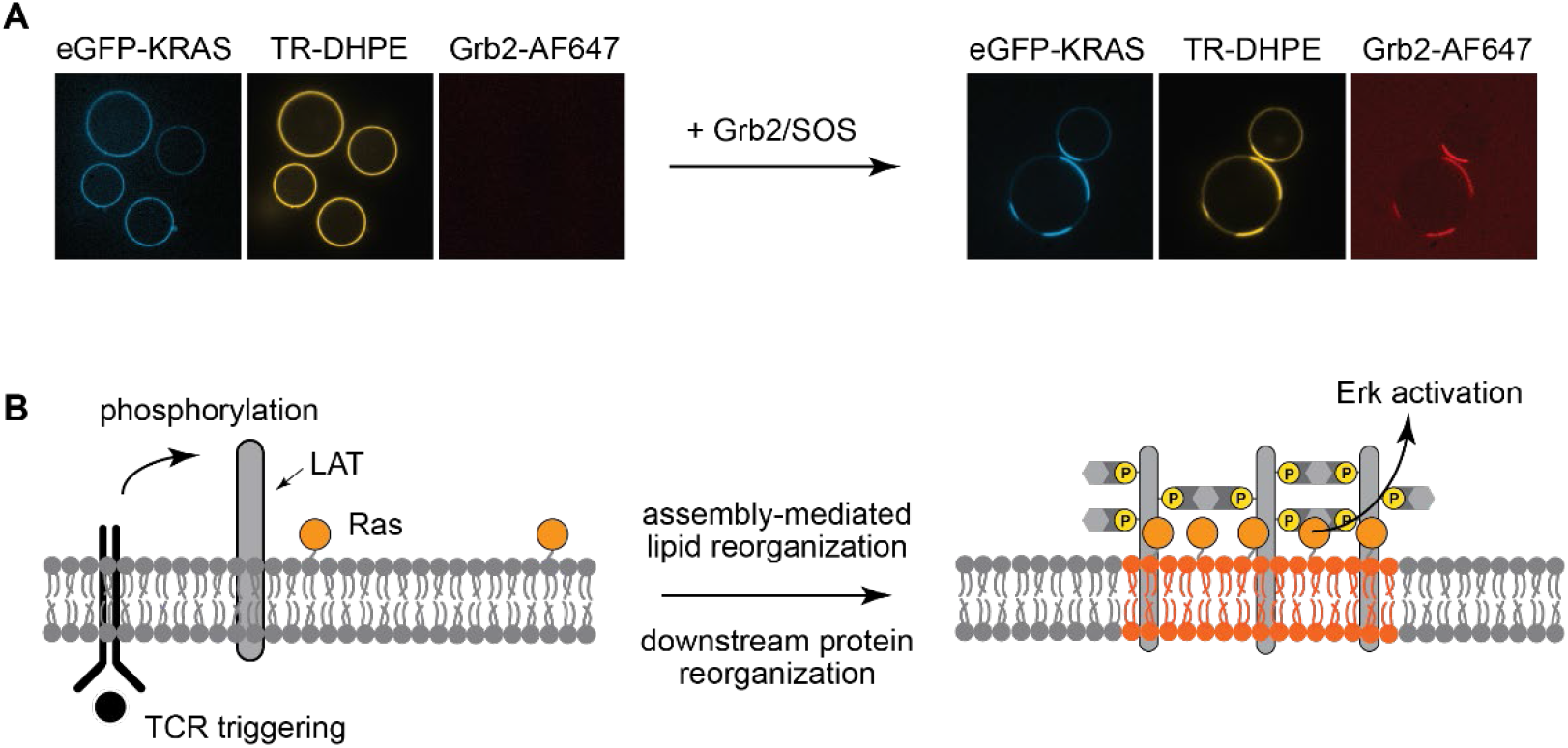
**(A)** The LAT:Grb2:SOS condensate on GUVs results in segregation of K-Ras into the *L*_d_ region with the condensate, suggesting that spatial organization mediated by protein assemblies can propagate downstream of the signaling pathway via lipids. **(B)** This lipid phase separation induced by protein organization may underlie lipid rafts seen in TCR clusters.

In summary, we have reconstituted the T-cell signaling condensate, LAT:Grb2:SOS, on vesicles capable of undergoing liquid-liquid miscibility phase transitions. We observed that the formation of protein condensate can drive the lipid phase transition under isothermal conditions, redistributing lipids in a signal-dependent manner. Furthermore, we have shown that K-Ras, which does not directly participate in the LAT:Grb2:SOS condensation, nonetheless colocalizes with the condensate through its sensitivity to the lipid environment. Lipid phase separation can also be induced by actin polymerization (72, 73) and lipid crosslinking by cholera toxin (74). Unique to the observations reported here, however, is that the LAT:Grb2:SOS protein condensation occurs immediately downstream of TCR activation, and as a direct result of ZAP70 kinase activation on triggered TCRs (28, 75). ZAP70 is a Syk Family Kinase that exhibits a distinctive substrate specificity, orthogonal to that of other kinases in the TCR signaling system, and strongly favors phosphorylation of the specific tyrosine residues on LAT that are involved in the LAT condensate (76). In this way, the LAT condensation phase transition is selectively controlled by TCR signaling.

The native LAT protein has been reported to exhibit a similar preference for the *L*_d_ lipid phase as the lipid-linked LAT in our experiments (38). However, in light of the significant number of other membrane-associated and transmembrane proteins in the cellular context of the LAT signaling condensate (59), we would refrain from extrapolating these results to predict specific details of the lipid phase associated with LAT in the natural physiological setting. The important point is that LAT condensation perturbs the underlying lipids and is capable of inducing lipid phase separation, now under the control of TCR signaling (Fig. 5). The LAT:Grb2:SOS protein condensate is not unique. Other two-dimensional condensates have been discovered, with their own signaling specificities (50, 51), and more are likely to emerge (e.g. with EGFR, which shares multivalent Grb2 and SOS interactions much like LAT). Such protein condensates on the membrane may play a broad role directly connecting receptor signaling activity with membrane lipid phase structure.

From a more physical perspective, a distinctive feature of the coupled protein-membrane system is that it exhibits phase transitions isothermally, and under control of competitive kinase-phosphatase reactions. At a single temperature and composition, the molecular interactions themselves change (as a function of LAT phosphorylation), and the phase state of the system follows. This differs from typical observations of lipid miscibility phase transitions, in which the molecular properties of the lipids are fixed, and other control parameters such as temperature potentiate the phase transition (16, 77–79). This control over LAT condensation through tyrosine phosphorylation not only enables the specific connection with cellular signaling systems, it also opens the door to various nonequilibrium chemical phenomena.

An example for such nonequilibrium phenomena can be found in a competitive lipid kinase-phosphatase reaction, which is similar to the tyrosine kinase-phosphatase competition governing LAT phosphorylation. The lipid kinase-phosphatase system has recently been observed to exhibit scale sensitivity in which the final outcome of the reaction depends on the size of the reaction system (e.g. a corralled lipid membrane in micron scales) (80). In this case, under identical concentrations of lipid kinases and phosphatases in solution, the membrane reaction system reaches a PIP_2_- or PIP_1_- (lipid kinase and phosphatase products, respectively) dominated state based on size and degree of confinement by the corralled membranes. Even partially confined membrane features, such as filopodia, are sufficient to flip the reaction outcome, and more elaborate pattern formations occur under different geometric restrictions. As with all kinase-phosphatase competitive cycles, this example is a dissipative process that continuously consumes ATP. The system is intrinsically out of equilibrium and the mechanism of this reaction scale sensitivity is rooted in nonequilibrium aspects of the kinetic system (81). The tyrosine kinase-phosphatase reactions upstream of the LAT condensate are qualitatively similar to the lipid kinase-phosphatase system mentioned above, albeit with even more complex feedback and regulatory couplings (82, 83). In the case of the LAT condensate, functionally critical properties such as nucleation threshold, size distribution, and growth-dispersion characteristics, are likely to be set by the kinase-phosphatase reactions controlling LAT phosphorylation. The LAT condensates, as well as any lipid phase structure they cause, will thus reflect the chemical states of the signaling system—including those arising from nonequilibrium processes—rather than equilibrium phase separation. At the present time, very little is known about physical characteristics of LAT condensates in living cells, leaving a wealth of opportunities for detailed studies of these systems. From a functional perspective, one may speculate that lipid miscibility phase separation in living cells is inextricably coupled to numerous specific protein assemblies and signaling processes, many of which are only beginning to gain visibility.

## Acknowledgments

This work was supported by the Novo Nordisk Foundation Challenge Program under the Center for Geometrically Engineered Cellular Systems. Additional support was provided by National Institutes of Health (NIH) National Cancer Institute (NCI) Physical Sciences in Oncology Network (PS-ON) project 1-U01CA202241 and by NIH grant P01 A1091580. We thank John-Paul Denson, William K. Gillette, and Andrew G. Stephen (NCI RAS Initiative, Frederick National Laboratory for Cancer Research) for the eGFP-K-Ras construct. We thank Young Kwang Lee, Hiu Yue Monatrice Lam, Shalini Low-Nam, and Emily C. Laubscher for assistance with the pilot experiments.

## Author Contribution

J. K. Chung, W. Y. C. Huang, and J. T. Groves conceived the research.

J. K. Chung, W. Y. C. Huang, C. B. Carbone, and L. M. Nocka performed experiments.

J. K. Chung, W. Y. C. Huang, C. B. Carbone, R. D. Vale, A. N. Parikh, and J. T. Groves analyzed data and interpreted results.

J. K. Chung, W. Y. C. Huang, and J. T. Groves wrote the manuscript.

All authors commented on the manuscript.

